# Loss of INPP5E affects photoreceptor outer segment membrane biogenesis in iPSC-derived human retinal organoids

**DOI:** 10.1101/2025.08.25.672060

**Authors:** Kae R. Whiting, Mariam G. Aslanyan, Tina Beyer, Katrin Dahlke, Karsten Boldt, Ronald Roepman

## Abstract

Mutations in the ciliary protein INPP5E, encoded by inositol polyphosphate-5-phosphatase E, can cause retinal degeneration as part of the ciliopathy Joubert Syndrome or non-syndromic retinitis pigmentosa (RP). INPP5E regulates the membrane makeup of the primary cilium, however its function in the specialized sensory photoreceptor cells of the human retina remain unclear. Here we utilize control and CRISPR/Cas9-generated *INPP5E* knock-out (*INPP5E^KD^*) human induced pluripotent stem cells (iPSCs) to generate retinal organoids (ROs). Through proteomic and immunofluorescence analysis we show that INPP5E plays an important role in early retinal development and photoreceptor progenitor cell differentiation. In mature ROs, INPP5E localizes to the connecting cilium of photoreceptors, and the loss of INPP5E leads to altered localization of ARL13B and Rhodopsin in mature photoreceptors. Furthermore, photoreceptor outer segment structure is affected, leading to elongated outer segment membranes in both cone and rod photoreceptors, suggesting an important role for INPP5E in photoreceptor outer segment membrane biogenesis. Together, these data underline the importance of INPP5E in retina development and photoreceptor structure and highlight the usability of retinal organoids to study protein function in a human context.

## Introduction

Photoreceptors in the retina carry highly specialized primary cilia that play a vital role in light detection and transduction. Primary cilia are small, hair-like, non-motile sensory organelles important during organismal development and tissue homeostasis (1), however the photoreceptor sensory cilium has a unique structure – including an extended transition zone called the connecting cilium (CC) and an outer segment (OS) filled with opsin-loaded outer segment discs – that allows the photoreceptor sensory cilium to function as a signal transducer during the phototransduction cascade (2). Pathogenic variants in genes encoding proteins that are important for photoreceptor function can cause inherited retinal degeneration (IRD) such as retinitis pigmentosa (RP), that affects around 1 in 4,000 individuals worldwide (3,4). RP is a progressive disorder characterized by the gradual loss of photoreceptors in the retina, initially presenting as night blindness followed by the progressive loss of vision (4,5). To date, upwards of 80 genes have been identified as RP causative genes (6), with manifestations of the disorder occurring as both non-syndromic, where RP is the only phenotype, and syndromic disorders that affect multiple organs alongside the retina. Ciliopathies are a class of developmental disorders caused by defects in the function of the cilium, with RP being a prominent phenotype amongst many ciliopathies (7).

One gene that has been found to cause both syndromic and non-syndromic RP when mutated is *Inositol polyphosphate-5-phosphatase E* (8,9), which encodes the protein INPP5E: a phosphatase important for regulating the phospholipid membrane of the primary cilium by cleaving the 5’ phosphate group from the phosphoinositide phosphatidylinositol 4,5-bisphosphate (PI(4,5)P_2_) to form phosphatidylinositol 4-phosphate (PI(4)P) (10–12). In doing so, INPP5E plays an important role in organismal development and tissue homeostasis by regulating signaling pathways (13–15), cell cycle entry (11,16,17), and neural patterning during early brain development (18). Pathogenic variants in *INPP5E* can lead to several disorders including MORM (mental retardation, truncal obesity, retinal dystrophy and micropenis) syndrome (OMIM #610156), Joubert Syndrome (JS, OMIM #213300), and non-syndromic retinitis pigmentosa (RP, OMIM#268000) (9,19,20).

To date, the role of INPP5E in photoreceptors has only been studied using mouse models. INPP5E localizes to the inner segment and connecting cilium of photoreceptors where it plays a role in protein trafficking and outer segment stability (21,22). However, mouse models do not always recapitulate the human phenotype of progressive retinal disorders, likely due to differences in the mouse and human genome (23) and the fact that mice are nocturnal animals. Therefore, they do not have a cone photoreceptor-rich fovea that facilitates daytime color vision in humans (24). Here we utilized human induced pluripotent stem cells (iPSC) with a CRISPR/Cas9-generated point mutation that renders the catalytic domain of INPP5E non-functional, leading to a functional knock-out of INPP5E (*INPP5E^KD^*; generated and characterized previously (18)) to generate retinal organoids (RO). We used these ROs subsequently to model the function of INPP5E in human retina development and photoreceptor maintenance.

The *INPP5E^KD^* ROs show a delay in photoreceptor precursor generation at early organoid timepoints, but this delay does not hinder the generation of more mature photoreceptors. We confirm the localization of INPP5E in the connecting cilia of wildtype photoreceptors but could not detect INPP5E in photoreceptors of *INPP5E^KD^* ROs. In mature ROs, there is an increase in the number of rod and cone photoreceptors and the structural morphology of both rod and cone photoreceptor outer segments (OS) is altered by the loss of INPP5E, leading to longer OSs compared to control organoids. Finally, we show that ARL13B and rhodopsin are mislocalized in *INPP5E^KD^* ROs. Together, this indicates a complex and crucial function for INPP5E during retina development and photoreceptor maintenance.

## Results

### *INPP5E^KD^* iPSCs differentiate into mature ROs

To study the role of INPP5E in human retina development and photoreceptor function, we differentiated both control and *INPP5E^KD^*iPSCs into ROs over the course of 230 days (Fig. 1A). Organoid morphology was tracked throughout development via brightfield microscopy, which showed proper retinal organoid development in both control and *INPP5E^KD^* ROs (Fig. 1B). Organoid area was measured at d35 and d230 based on brightfield images, where there was no difference between control and mutant organoids (Fig. 1D). Through immunofluorescence staining of DAPI of cryopreserved and sectioned organoids, the retinal progenitor and retinal cell containing ONL could be visualized (Fig. 1C) and quantified (Fig. 1E). No difference in ONL length of developing (d35) and mature (d230) organoids was seen. Interestingly, though, there was a difference between the number of nuclei in the ONL of control and *INPP5E^KD^* organoids (Fig. 1F). At d35, there was a slight decrease in the number of nuclei per 50µm^2^ in *INPP5E^KD^* organoids (53.5 ± 9.2 nuclei/50µm^2^) compared to the control (69.3 ± 6.4 nuclei/50µm^2^). However, at d230, the number of nuclei per 50µm^2^ significantly increased in *INPP5E^KD^* ROs compared to control (control: 74.8 ± 2.9 nuclei/50µm^2^; *INPP5E^KD^*: 100.3 ± 14.6 nuclei/50µm^2^).

**Figure 1.**
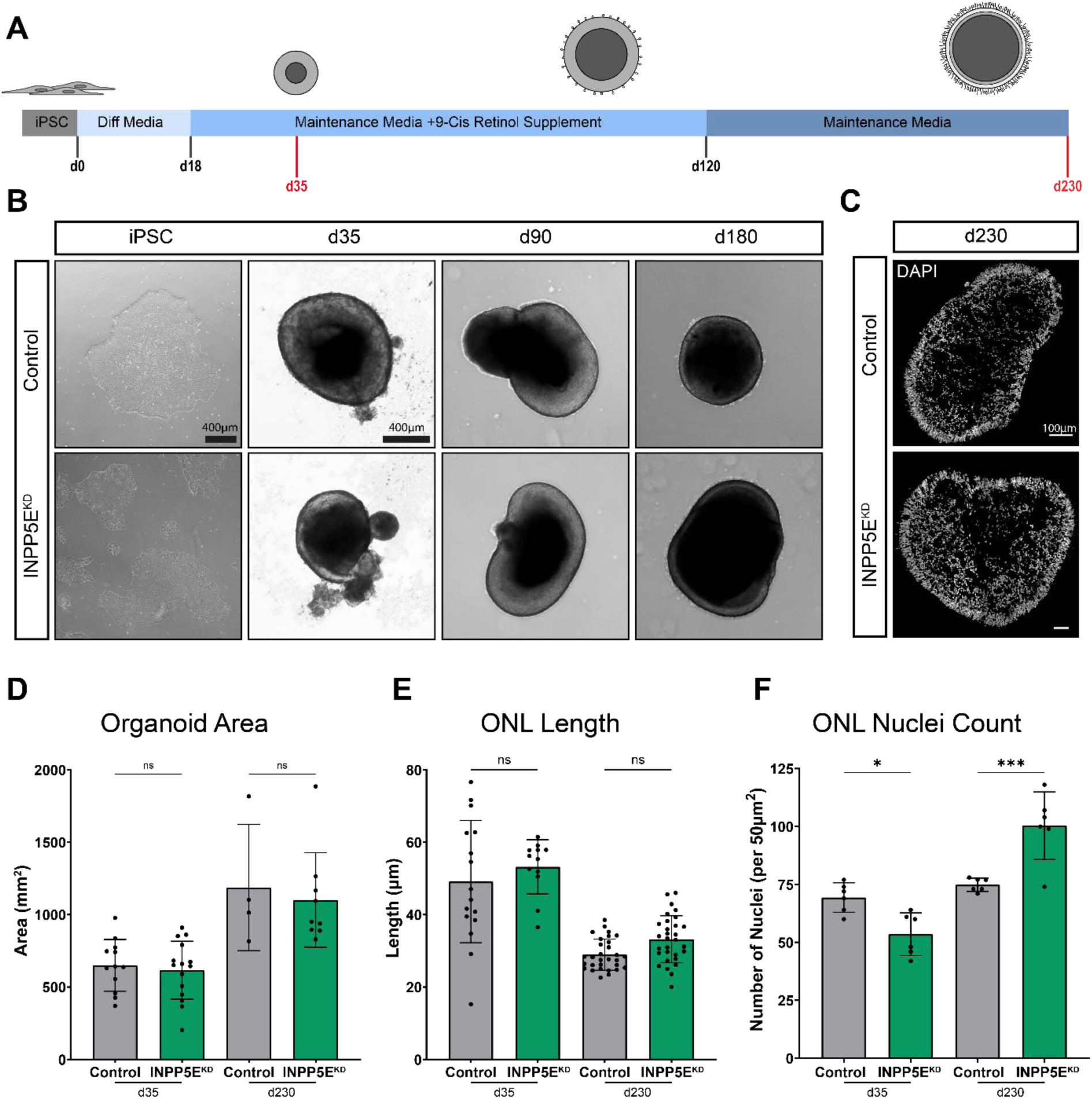
Differentiation of iPSCs to retinal organoids. (A) Protocol overview to differentiate iPSCs to RO. Makeup of Differentiation (Diff) and Maintenance Media can be found in methods. Organoids were collected at d35 and d230 for IF and proteomic analysis. (B) Brightfield images of organoids through development of control and INPP5E^KD^ organoids show no major morphological changes (Scale bar: 400µm). (C) IF of mature organoids at d230 stained for DAPI (Scale bar: 100µm). (D-F) Quantification of basic retinal organoid morphology at early (d35) and late (d230) stages of development as (D) retinal organoid area, (E) outer nuclear layer (ONL) length, and (F) quantity of DAPI in the ONL. (* p=0.0371, *** p=0.0007)

Further, we used immunofluorescence analysis of the photoreceptor precursor markers RCVN and CRX to confirm the differentiation towards retinal organoids and to determine if there are any phenotypic differences between control and knock-out organoids in early development (Fig. 2A). At early stages of control retinal organoid development (d35), the photoreceptor precursor markers Recoverin (RCVN) and CRX are initially expressed. *INPP5E^KD^* organoids showed a delay in photoreceptor precursor expression. Compared to control, *INPP5E^KD^*ROs had a slight, but not significant, decrease in CRX positive cells (control: 20.42%; *INPP5E^KD^*: 11.92%) and a significant change in RCVN expression in the ONL (control: 126.5 ± 36 A.U.; *INPP5E^KD^*: 64.28 ± 43 A.U.). (Fig. 2B,C).

**Figure 2.**
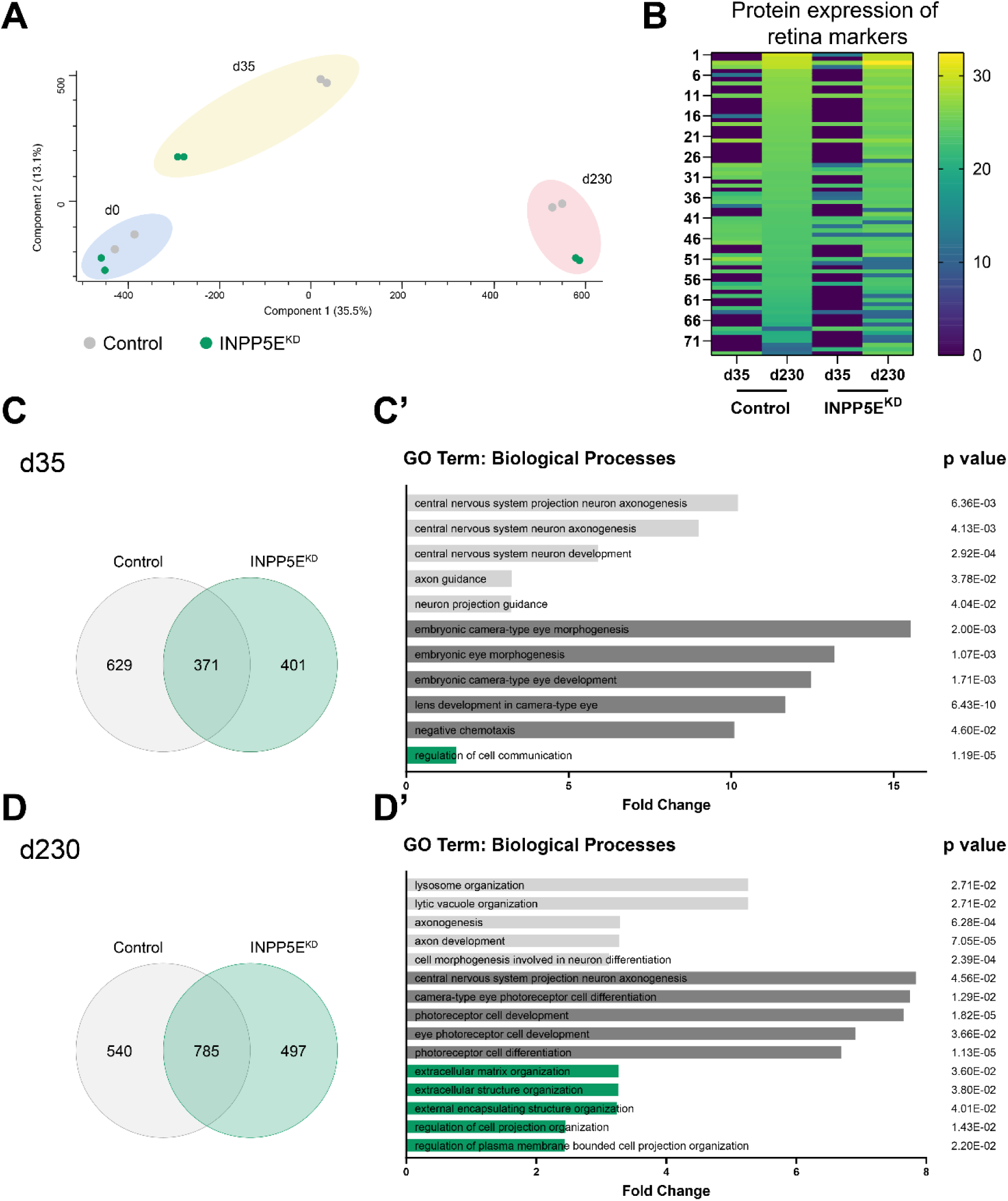
Bulk proteomics of organoids confirms retinal fate. (A) PCA plot of d0, d35, and d230 organoids show a trajectory of development as organoids differentiate further from iPSCs. (B) Protein expression of retina markers increases from d35 to d230 as organoids develop in both control and INPP5E^KD^ ROs. Comparison of enriched protein datasets at d35 (C) and d230 (D) of organoid development show high protein overlap at d230. (C’, D’) Top GO terms for biological processes of enriched proteins at d35 and d230. GO terms for proteins enriched in control organoids are light grey, in only INPP5E^KD^ organoids in green, and in both datasets in dark grey.

### Proteomics of retinal organoids confirms retinal fate

To confirm retinal differentiation of iPSCs to ROs, bulk proteomics was performed on undifferentiated iPSCs (d0), early stage ROs (d35) and mature ROs (d230). PCA revealed distinct clusters based on differentiation stage, demonstrating that mature ROs cluster further apart from iPSCs (Fig. 3A). Additionally, the expression of retinal specific markers – such as ARR3, RHO, and PRPH2 – markedly increased from d35 to d230 (Fig. 3B).

**Figure 3.**
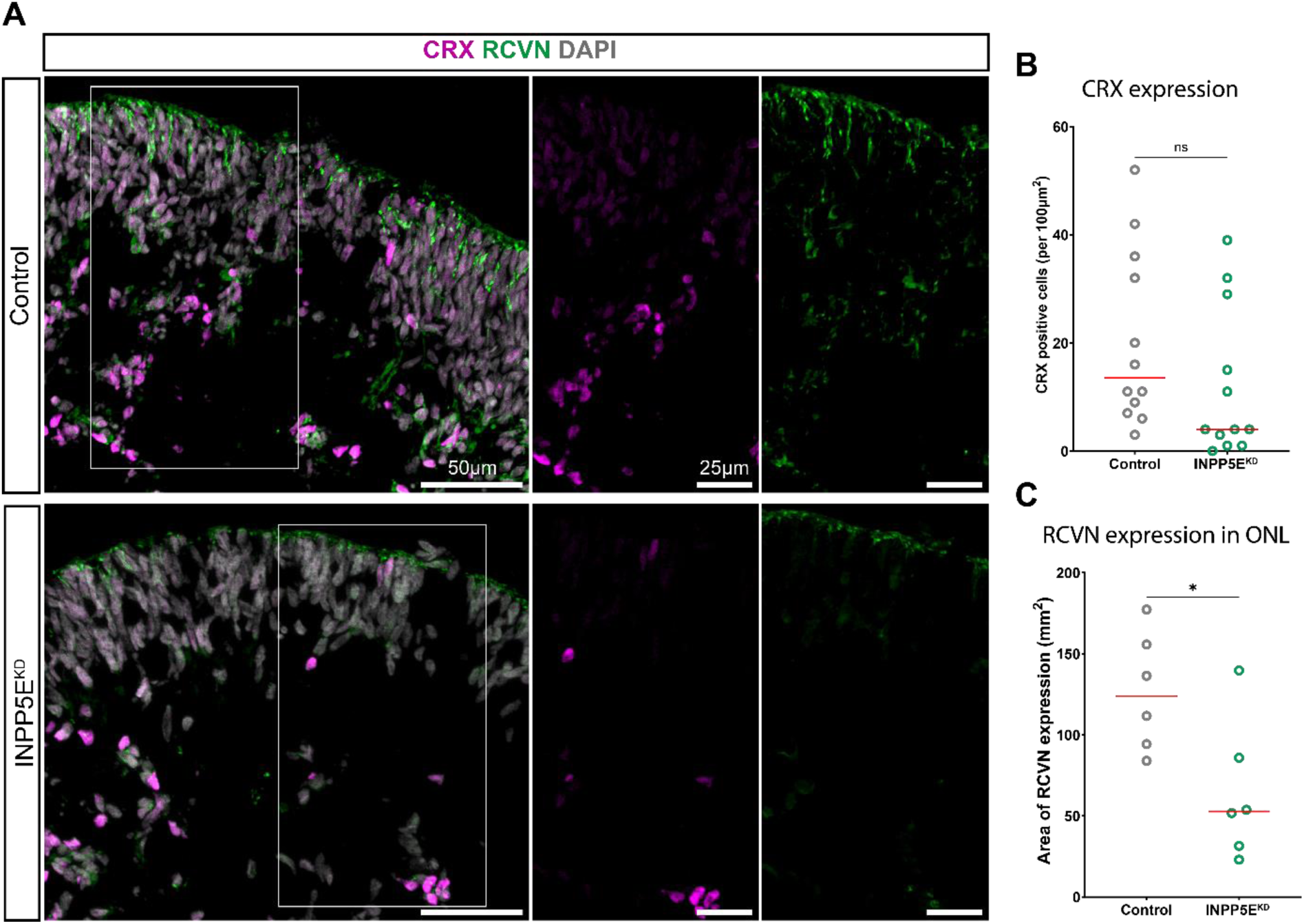
INPP5E^KD^ organoids have delayed expression of photoreceptor precursor markers. (A) Representative immunofluorescence of retinal precursor markers CRX (magenta) and RCVN (green) at d35 of differentiation (Scale bar: 50µm and 25µm) (B-C) Quantification of photoreceptor precursor markers as the percent CRX positive cells (C) and Area of recoverin (RCVN) expression in the ONL (D). (* p<0.05, **** p<0.0001).

To better understand the mechanism of INPP5E within the ROs, Gene Ontology (GO) enrichment analysis for biological processes (BP) was performed. Proteins were classified into three groups: proteins enriched in only control ROs (light grey), proteins enriched in only INPP5E^KD^ ROs (green), and proteins enriched in both datasets (dark grey) (Fig. 3C,D). During early RO differentiation (d35) several BP terms for embryonic eye development were enriched in both control and *INPP5E^KD^* ROs. BP terms for axon guidance and axonogenesis were highly enriched in control organoids, while in *INPP5E^KD^* ROs regulation of cell communication was the only significant term (Fig. 3C’). In mature retinal organoids (d230), BP terms for photoreceptor cell development were highly enriched in both datasets. However, there was a difference in GO terms identified for control and *INPP5E^KD^*ROs: control ROs were enriched for axonogenesis and lysosome organization while *INPP5E^KD^* ROs were enriched for cell projection organization and extracellular matrix organization (Fig. 3D’).

### Cilia morphology and INPP5E expression is affected in *INPP5E^KD^*ROs

In most cell models, ARL13B regulates INPP5E localization at the ciliary membrane of the primary cilium, where the two co-localize (35). In the retina, INPP5E localizes to the inner segment and connecting cilium of photoreceptors in mouse, non-human primate, and the mature human macula (22). Moreover, previous studies have shown that defects in INPP5E, due to either patient-specific mutations or CRISPR/Cas9 generated knock-outs, lead to both the mislocalization of INPP5E to the cilium and shorter cilia (8,18). Here, we used immunofluorescence of INPP5E and ARL13B to determine the localization of INPP5E in developing and mature ROs. In control organoids, INPP5E colocalized with ARL13B along the primary cilium of both d35 ROs (2627 ± 1669 A.U.) and mature d230 retinal organoids (2629 ± 1505 A.U.). As expected, in *INPP5E^KD^*organoids, INPP5E expression was significantly decreased at both d35 and d230 of RO development (d35: 948.1±605 A.U.; d230358.7±496.5 A.U.) (Fig. 4E).

**Figure 4.**
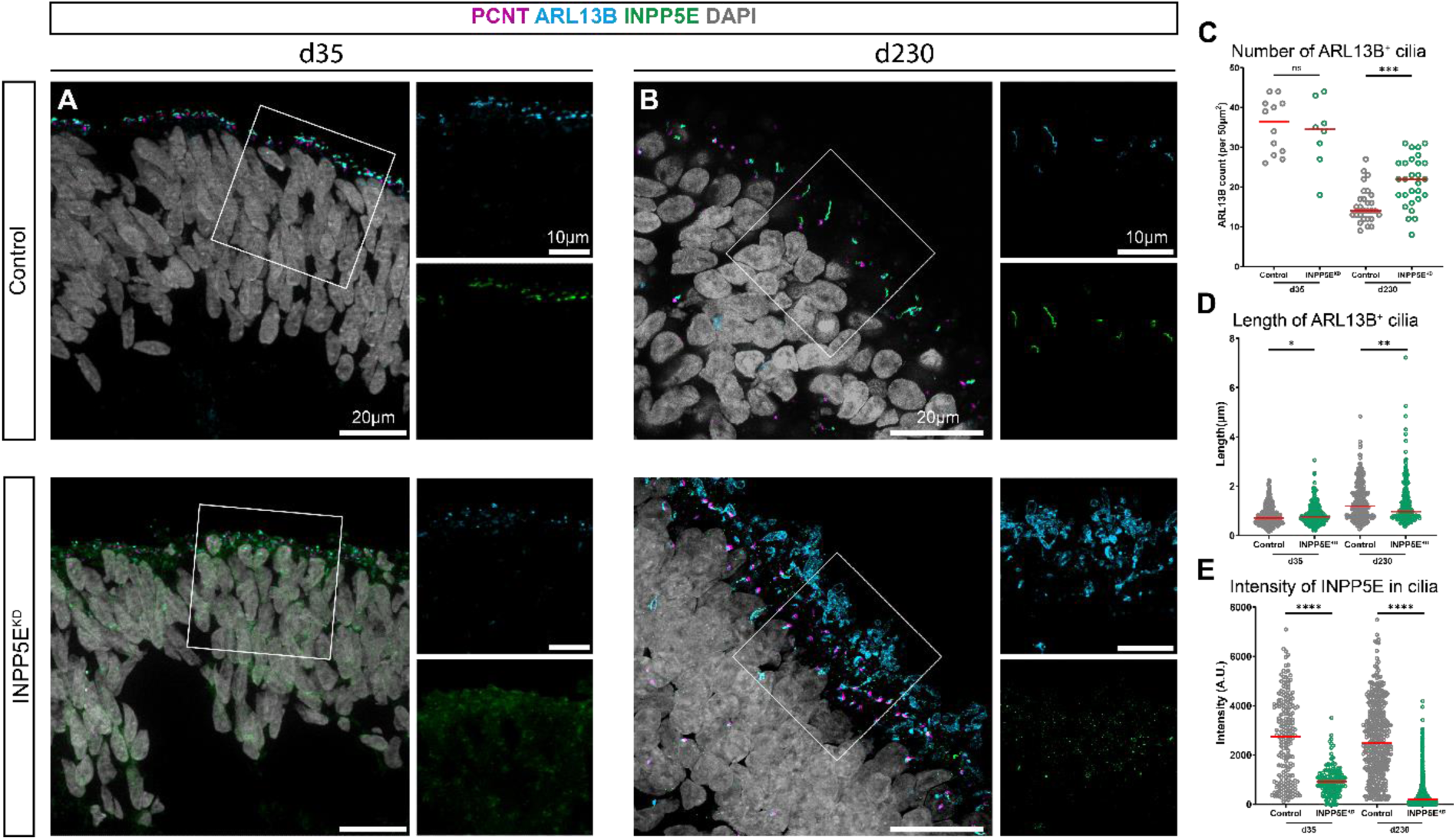
Structural changes of primary and connecting cilia in INPP5E^KD^ organoids. (A-B) Representative immunofluorescent images of d35 (A) and d230 (B) ROs stained for the basal body marker PCNT (magenta), the ciliary membrane marker ARL13B (blue), DAPI (grey) and INPP5E (green). (E-G) Quantification of ciliary morphology and INPP5E localization as the number of ARL13B^+^ cilia per 50µm^2^ (C), length of cilia (µm) (D), and intensity of INPP5E expression in the cilia (A.U.) (G). (* p = 0.0139, ** p = 0.0083, *** p = 0.0001, **** p <0.0001).

Next, to determine if the loss of INPP5E affects ciliary morphology during RO differentiation we utilized immunofluorescence to visualize the ciliary markers pericentrin (PCNT) and ARL13B in d35 and d230 ROs (Fig. 4A,B). In control ROs both the number and structure of cilia in control retinal organoids changed over the course of differentiation. At d35, there were more cilia than at d230 (d35: 35±6.8 ARL13B^+^ cilia per 50µm^2^; d230: 15±4.4 ARL13B^+^ cilia per 50µm^2^) (Fig. 4C) while the length of cilia increases as the organoids develop and mature (d35: 0.797±0.38 µm; d230: 1.367±0.73 µm) (Fig. 4D).

*INPP5E^KD^* organoids show slight defects in ciliary morphology during early development, with no significant differences in the number of cilia at d35 (33±8.4 ARL13B^+^ cilia per 50µm^2^) (Fig. 4C) and a minor increase in cilia length compared to control (0.87±0.41 µm) (Fig. 4D). However, at d230 the number of ARL13B^+^ cilia significantly increases compared to control (21 ± 6.2 ARL13B^+^ cilia per 50 µm^2^) while the length of the cilia decreases (1.27 ± 0.88 µm) (Fig. 4C,D).

To determine if other ciliary sub-compartments were affected, we used immunofluorescence of the basal body marker PCNT and the ciliary TZ marker NPHP1. Expression of PCNT and NPHP1 was present at the basal region of the CC and no phenotypic alterations were apparent between control and *INPP5E^KD^*ROs (Supp Fig. 1).

### Photoreceptors of INPP5E^KD^ ROs have significant morphological defects

To better understand the role of INPP5E in the development and maintenance of mature photoreceptors, we utilized immunofluorescence to visualize rod and cone photoreceptors in mature ROs, using Rhodopsin (RHO) and Opsin R/G respectively (Fig. 5A). In control ROs, there are 100 RHO^+^ photoreceptors and 47 Opsin^+^ photoreceptors per mm, giving a 69/31% rod/cone photoreceptor ratio. In *INPP5E^KD^*ROs, the ratio of rod/cone photoreceptors was not altered (71/29% rod/cone), however there was an increase in the total number of both the rod and cone photoreceptors compared to control ROs (150 Rho^+^ photoreceptors per mm; 57 Opsin^+^ photoreceptors per mm) (Fig. 5B-D). Moreover, a significant increase in the length of the outer segments in both cone (control: 5.87 ± 2.6 µm; *INPP5E^KD^*: 7.13 ± 4.5 µm) and rod (control: 6.71 ± 3.5 µm; *INPP5E^KD^*10.90 ± 5.9 µm) photoreceptors was observed in the INPP5E^KD^ ROs (Fig. 5E,F).

**Figure 5.**
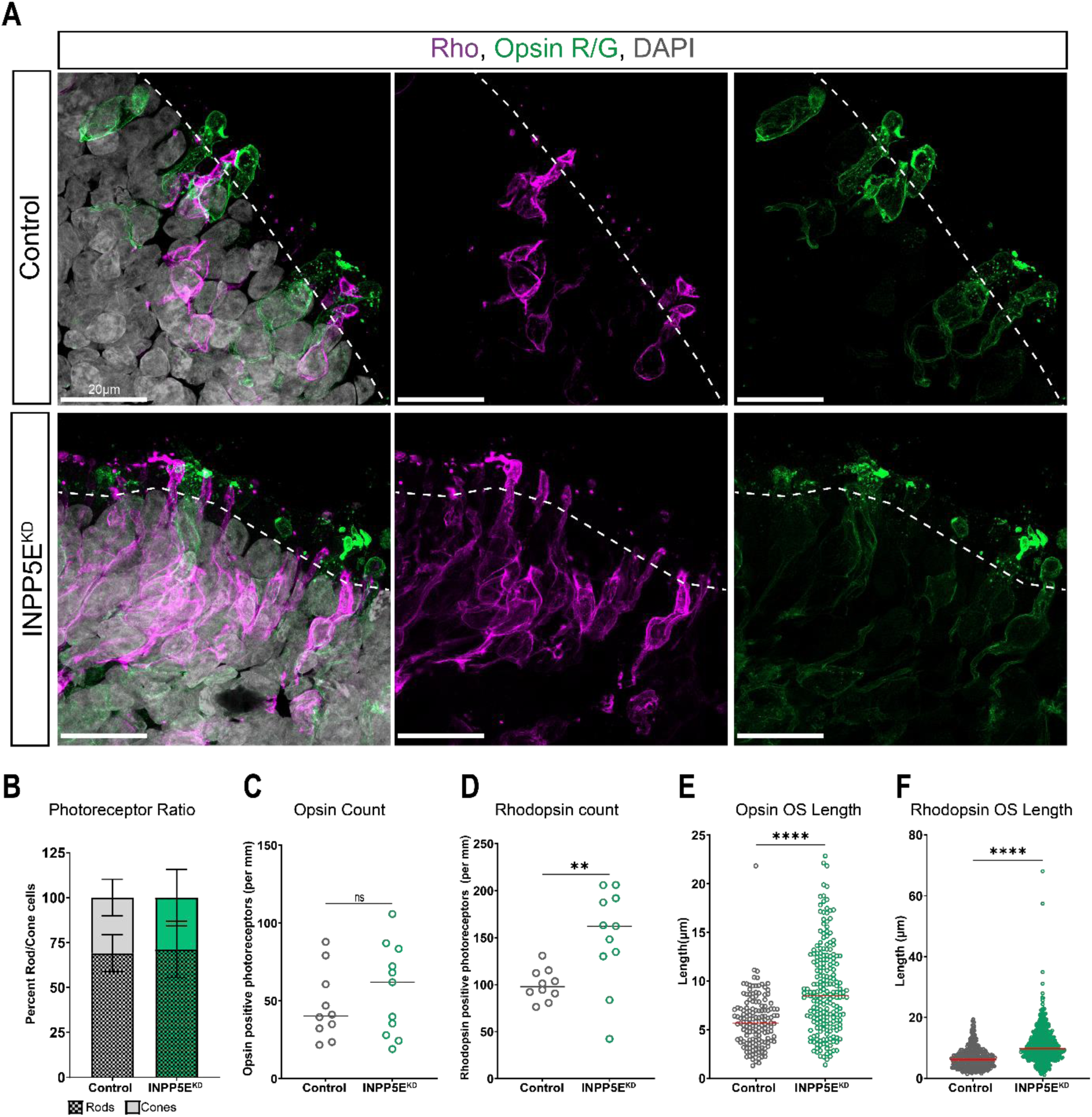
Photoreceptor morphology is altered in INPP5E^KD^ organoids. (A) representative images of rod and cone photoreceptors in d230 organoids, stained for RHO (magenta), Opsin R/G (green), and DAPI (grey). Dashed line represent outer limiting membrane of organoids (scale: 20µm). (B) Quantification of rod/cone ratio in control and INPP5E^KD^ organoids as the percent RHO positive and opsin positive photoreceptors. (C-D) Quantification of photoreceptor development as the number of opsin positive (C) or rhodopsin positive (D) photoreceptors per mm. (E-F) Comparison of photoreceptor morphology as the length of the photoreceptor outer segment (OS) for opsin positive cone photoreceptors (E) and rhodopsin positive rod photoreceptors (F). (** p=0.0013; **** p<0.0001).

### Altered actin and tubulin regulation in *INPP5E^KD^* organoids

Previous work has suggested a role for INPP5E in regulating actin in mouse photoreceptors (21). Therefore, we speculated that the same could be true in retinal organoids. We utilized phalloidin to visualize filamentous actin (F-actin) and determine if INPP5E plays a role in regulating the actin network in developing photoreceptors. In control ROs, phalloidin was expressed at the outer limiting membrane (Fig. 6A), however small extensions of phalloidin were visible extending into the CC of photoreceptors. In *INPP5E^KD^* ROs phalloidin co-localized with the outer limiting membrane, however it extended further into the CC than in control ROs (control: 2.37 µm; INPP5E^KD^ 3.37 µm) (Fig. 6C).

**Figure 6.**
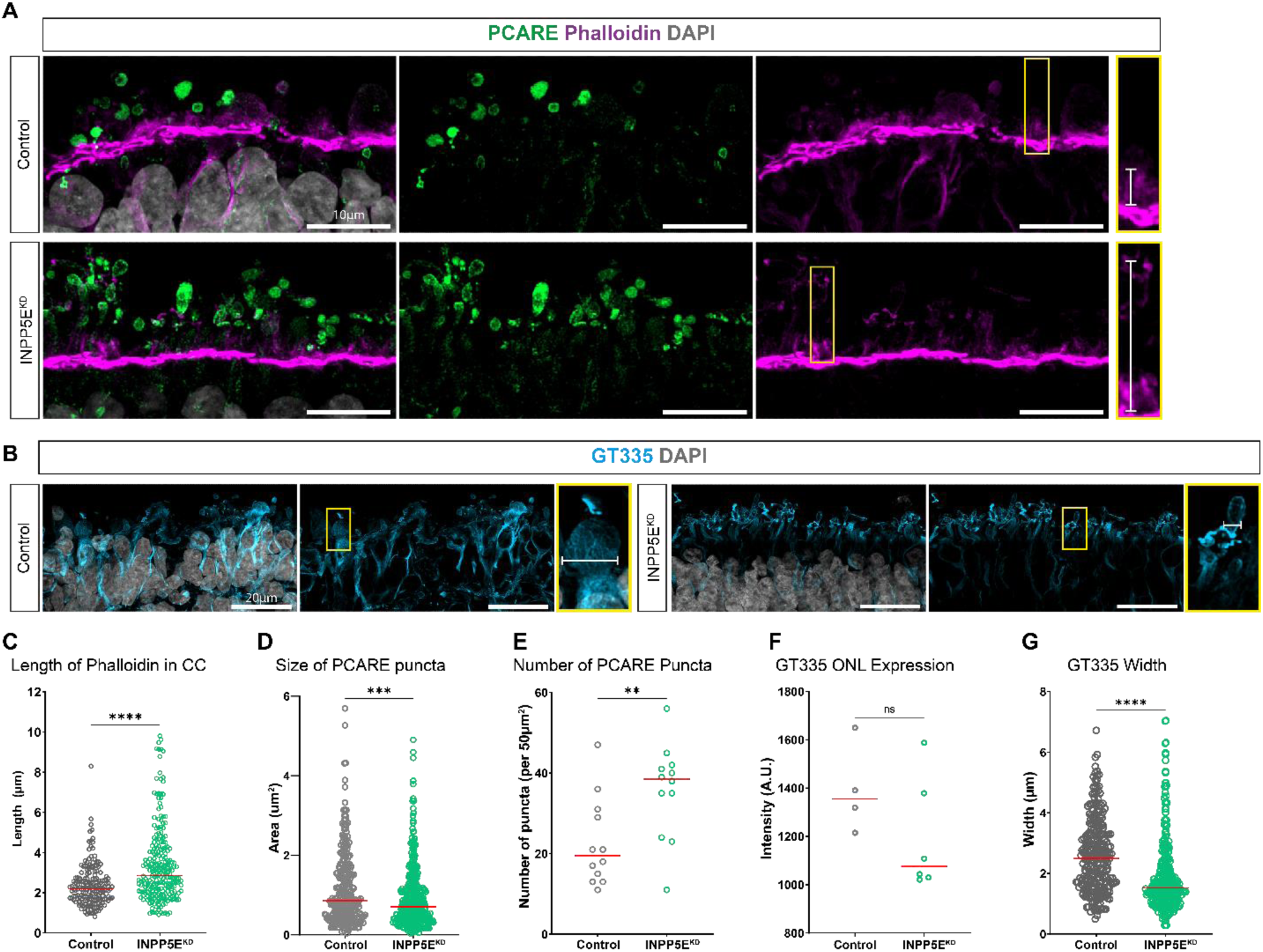
Actin and tubulin dysregulation in d230 INPP5E^KD^ organoids. (A) Representative immunofluorescence of f-actin (phalloidin, magenta), the actin regulator PCARE (green), and DAPI (grey) in d230 control and INPP5E^KD^ ROs. Yellow cutouts show examples of phalloidin extension into the photoreceptor connecting cilium (Scale bar: 10 µm). (B) Representative images of polyglutamylated tubulin (GT335, blue) and DAPI (grey) expression in d230 ROs. Cutout shows example of GT335 tubulin bundle measurement. (Scale: 20 µm). Quantification of actin and actin regulators as the length of phalloidin extending into the connecting cilium (C), the size of PCARE puncta in photoreceptor OS (D), and the number of PCARE puncta per 50µm^2^ (E). Quantification of GT335 expression was performed as the intensity of GT335 expression in the ONL (F) and the spread of GT335 tubulin bundles in photoreceptor axonemes (G). (** p = 0.0099; *** p = 0.0001; **** p<0.0001)

To gain further insight into the possible dysregulation of the actin protein network, we performed a co-localization experiment with the photoreceptor actin regulator PCARE, which is expressed at the base of the OS and plays an important role in the initiation of new outer segment disc formation (36,37). The number of PCARE puncta were increased in *INPP5E^KD^* organoids (control: 22.7 puncta per 50µm^2^; INPP5E^KD^: 35.8 puncta per 50 µm^2^), as seen previously with ARL13B and RHO. However the area of the PCARE puncta was decreased in *INPP5E^KD^* organoids (0.91 µm2) compared to control (1.16 µm^2^) (Fig. 6D,E).

We next sought to determine if the loss of INPP5E affected photoreceptor microtubules. To do so, we used the polyglutamylated tubulin marker GT335. The number of GT335^+^ photoreceptors increased in *INPP5E^KD^* ROs compared to control at d230 (control: 31.8 ± 6.4; *INPP5E^KD^* 42.9 ± 5.7), while the intensity of GT335 expression in the ONL was slightly, but not significantly, decreased in *INPP5E^KD^*compared to control ROs (control: 1394 A.U.; *INPP5E^KD^*: 1195 A.U.) (Fig. 6B,F). However, interestingly, the loss of INPP5E led to an altered morphology of GT335^+^ tubulin in the photoreceptors, causing the spread of tubulin bundles in the axoneme, measured by the width of GT335 staining, to decrease compared to control (control: 2.62 µm; *INPP5E^KD^*: 1.88µm) (Fig. 6G).

### Rhodopsin and ARL13B localization are altered in *INPP5E^KD^* ROs

Previous studies have demonstrated a defect in rhodopsin trafficking in INPP5E mutant mice (21,22), with further studies showing that retention of rhodopsin in the outer nuclear layer of the retina being indicative for several known IRDs (21,38,39). In line with these studies, immunofluorescence of RHO showed an increase of RHO expression in the ONL of *INPP5E^KD^* ROs compared to control (control: 375±103.6 A.U.; *INPP5E^KD^*: 708 ± 260.6 A.U.) (Fig. 7A,B). We also analyzed opsin R/G localization in the ONL, but saw no change in localization, suggesting that the trafficking defect is rhodopsin specific (Supp Fig. 2).

**Figure 7.**
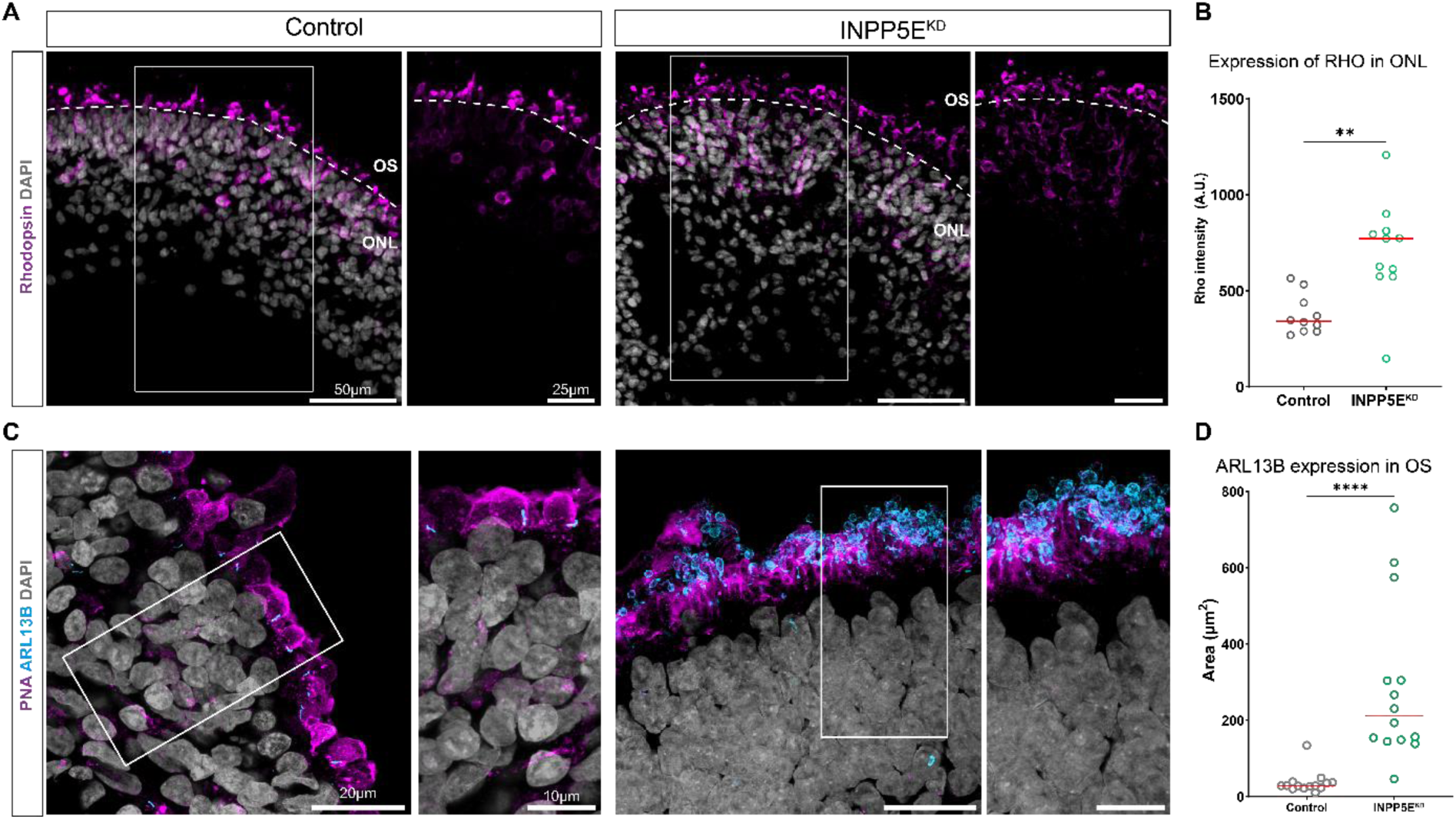
Rhodopsin and ARL13B trafficking is altered in d230 INPP5E^KD^ organoids. (A) Representative immunofluorescence of rhodopsin (RHO) and DAPI (grey) in the OS and ONL. The white dashed shows the separation of the ONL and OS of the ROs (scale: 50 µm and 25 µm). (B) Quantification of expression of RHO in the ONL, measured as the normalized intensity (A.U.). (C) Representative images of ARL13B (blue), PNA (magenta), and DAPI (grey) (scale: 20 µm and 10 µm). (D) Quantification of ARL13B expression in PNA^+^ OS, measured as the total area (µm^2^). (** p = 0.0013, **** p<0.0001).

Interestingly, in *INPP5E^KD^* RO we saw an increase of ARL13B expression outside the photoreceptor CC, suggesting a change in the protein localization. To confirm this, we visualized ARL13B with peanut agglutinin (PNA), which stains the outer segment membrane of photoreceptor cells (40) (Fig. 7C). In control organoids, the area of ARL13B expression in the PNA^+^ OS was 36.6 µm^2^, which was significantly increased to 288.2 µm^2^ in *INPP5E^KD^* organoids.

## Discussion

Pathogenic variants that cause dysfunction in *INPP5E* lead to both syndromic and non-syndromic RP in patients (8,9), suggesting an important role for INPP5E in retinal function. While recent studies have utilized conditional mouse models that selectively silence INPP5E in the retina or photoreceptors (21,22), mouse models do not always recapitulate the human phenotype when modeling retinal dysfunction (23,41). Here we utilized retinal organoids to study the role of INPP5E in retina development and photoreceptor maintenance in a human model system. Both control and *INPP5E^KD^* organoids differentiate successfully with no major differences in the general structure of the retinal organoids, suggesting that INPP5E is not critical for retinal development.

In mature ROs, photoreceptors develop as part of a so-called “brush border” along the outer edge of the organoid, emerging as early as d150 and maturing by d230. These photoreceptors express both rod and cone-specific proteins including rhodopsin and opsin, respectively. The number of both rod and cone photoreceptors in *INPP5E^KD^* ROs were increased compared to control organoids, however the outer segment membranes of photoreceptors in *INPP5E^KD^* organoids were significantly longer than control ROs. Previously published work using conditional knock-out mouse models of INPP5E showed that the loss of INPP5E led to a decrease in outer segment length due to a decrease in OS disc stability – opposite to what was seen previously in the *INPP5E^KD^* ROs (21,22). This opposing result is likely due to the differences in the structure of outer segments in *in vivo* mouse models versus the *in vitro* retinal organoids. Typically, outer segments are filled with highly organized outer segment discs, responsible for light capture to initiate phototransduction. While retinal organoids are able to generate short outer segments with rudimentary outer segment discs, they lack the structure and organization of *in vivo* photoreceptors. This is likely due to the lack of the retinal pigment epithelium (RPE) cell layer, which provides both structural and functional support to the photoreceptor OS (42).

It has been hypothesized that alongside outer segment stability, INPP5E also plays a role in regulating protein trafficking between the inner and outer segment (21). Here, mature *INPP5E^KD^* organoids had accumulation of rhodopsin in the ONL, a phenotype seen previously in multiple RP-associated models including the INPP5E mouse knock-out model (21,38), validating this as a phenotypic consequence in humans. Similar to the INPP5E mouse model (21), we could demonstrate that INPP5E depletion causes defects in the actin cytoskeleton, but we also did identify differences in the expression of the photoreceptor actin regulator PCARE. This matches the known role of INPP5E in actin regulation. The extended cc region that is covered by actin could be indicative of a disrupted balance in actin dynamics, potentially due to enhanced PI(3,4,5)P_3_and PI(4,5)P_2_ levels that are not converted due to absence of INPP5E. Raised levels of PI(3,4,5)P_3_ promote AKT phosphorylation, altering actin dynamics, and raised levels of PI(4,5)P_2_ promotes actin nucleation through interaction with the WAVE complex. A disturbed balance of these processes may underly our observations of the extended OS as well as increased levels but altered size of PCARE puncta. The altered tubulin dynamics could then be a result of the disturbed cytoskeletal processes, that are intertwined at the base of the outer segments to facilitate proper outer segment membrane biogenesis. The fact that somewhat different observations were done in the INPP5E knock-out mouse model, where reduced phalloidin staining but no alterations in WAVE complex was observed (21), could indicate that the early imbalance in actin dynamics that we observe in the retinal organoids triggers developmental defects that are expressed in altered stoichiometry in the actin dynamics network at postnatal stages. Although tempting, such hypotheses should be made with caution, as also species differences could be at play.

Further, mature *INPP5E^KD^*organoids showed mislocalization of ARL13B, which localizes ectopically to the outer segments of INPP5E^KD^ photoreceptors, strengthening the hypothesis that INPP5E regulates protein trafficking within the specialized photoreceptor cilium.

This change in ARL13B localization in *INPP5E^KD^* photoreceptors might help explain the elongation of photoreceptor outer segments. ARL13B is a regulatory GTPase found at the membrane of the primary cilium, with previous work showing that ARL13B plays a role in regulating ciliary length, where increased ARL13B expression results in ciliary elongation (43,44). The loss of INPP5E function leading to ARL13B mislocalization could be the cause for changes in membrane biogenesis and thus longer OS in the knock-out photoreceptors.

Interestingly, ARL13B mislocalization appears only in later phases of organoid differentiation as ARL13B localization and expression were unaffected in the cilia of d35 *INPP5E^KD^* organoids, possibly alluding to a different function of INPP5E during early retinal differentiation. In addition, INPP5E loss led to a delay in photoreceptor precursor cells in early stage ROs, such as CRX expressing cells, though this delay was rescued by the time organoids reached maturation at d230.

A possible scenario for this distinct contribution during early RO development comes from studies of INPP5E function in early brain development. The authors of the study used the same *INPP5E^KD^* and isogenic control iPSC lines to generate cerebral organoids, where they demonstrate that the loss of INPP5E leads to an increase in sonic hedgehog (SHH) signaling, thus playing an important role in dorsal ventral patterning during brain development (18). In the retina, SHH plays a role in regulating cell proliferation and differentiation during retinogenesis – with the activation of SHH leading to faster division of proliferative cells (45). Therefore, it is possible that the loss of INPP5E in early retinal organoid development leads to increase in SHH signaling, thus increasing the pool of proliferative cells that would go on to form retinal precursor and photoreceptor cells – explaining the increase in DAPI^+^ cells, the increase of total rod and cone photoreceptor cells in mature ROs, and the increase in PCARE and NPHP1 puncta within these photoreceptors. Future work to assess the role of SHH signaling during retinogenesis will help elucidate this hypothesis further.

To conclude, we generated retinal organoids from iPSCs containing a knock-out of INPP5E, which we used to expand upon previous knowledge of the function of INPP5E in both developing and mature photoreceptors. This indicates a complex function of INPP5E in the retina, both during early retinal differentiation and in mature photoreceptors. Further, we show that retinal organoids can be a useful model system to study INPP5E-derived retinal degeneration. In the future, this model system can be used to study potential therapeutic approaches to rescue the phenotype caused by the loss of INPP5E.

## Methods

### iPSC-derived retinal organoid differentiation

Human induced pluripotent stem cells (iPSCs) were cultured on Matrigel (Corning, 734-0269) coated 6-well plates in mTesR1 media (STEMCELL Technologies, 85850) containing 1% penicillin-streptomycin (Pen/Strep. Sigma-Aldrich, P4333). Cells were passaged with 0.5 mM EDTA (Invitrogen, 11568896) and maintained at 37°C with 5% CO_2_. CRISPR/Cas9 generated INPP5E knock-out iPSCs, containing a homozygous point mutation (D477N/D477N, INPP5E^KD^) and their corresponding isogenic control line, characterized previously (18) were a kind gift from Dr. Thomas Theil (University of Edinburgh, UK).

iPSCs were differentiated into retinal organoids using a previously established protocol (25,26). Briefly, iPSCs at 75-80% confluency was dissociated as single cells using Accutase (Gibco, A1110501) and seeded at 7,000 cells per well in ultra-low adhesion U-Bottom 96-well plates (Corning, 7007) in mTesR1 media containing 10 µM Y-27632 (Rock inhibitor; Sigma-Aldrich, Y0503). After 48 hrs (day 0 of differentiation), 200 µm differentiation medium (41% IMDM (Sigma, I3390), 41% HAM’s F12 (Sigma, N6658), 15% KOSR (Gibco, 10828028) 1% GlutaMAX (Gibco 35050061), 1% Lipids (Gibco, 11905031), 1% Pen/Strep, and 0.2% 1-Thioglycerol (Sigma-Aldrich, M6145)) was added to each well. Every other day, organoids were subjected to half media changes, and on day 6 of differentiations a 2.25 nM BMP4 (Sigma, SRP3016) pulse was added. At d18, retinal organoids were moved to ultra-low adhesion 6-well plates (Corning, 3471), and the medium was changed to Maintenance medium containing 9-cis-retinol (DMEM/F12 (Gibco, 31330038), 10% FBS, 1% N2 (Gibco, 17502048), 1% Pen/Strep, 1% GlutaMAX, 01. mM Taurine (Sigma Aldrich, T0625), 0.25ug/mL Amphotericin B (Gibco, 15290018), and 0.5 µM 9-cis-Retinol (Sigma-Aldrich, R5754)). Media was changed every 3-4 days, and 9-cis-retinol was removed after 120 days of differentiation.

Brightfield images of retinal organoid differentiation were captured at multiple timepoints throughout the differentiation using the EVOS XL Core Cell Imaging System (Invitrogen).

### Immunofluorescence

Two differentiations were completed per cell line and 3-4 ROs were collected at day 35 (d35) and day 230 (d230) per differentiation. ROs were washed in PBS and fixed in 4% PFA at room temperature (RT) for 20 minutes and cryoprotected in 30% sucrose overnight at 4°C. ROs were then embedded in OCT compound (Sakura Finetek Europe, 25322-68-3), and slowly frozen on dry ice. Cryosections were collected at 12µm thick and stored at −20°C until used for immunofluorescence.

In short, sections were washed briefly in PBS to remove OCT and incubated in blocking buffer (0.3% Triton-X, 2.5% natural goat serum (NGS) in PBS) for 1 hour at RT. Slides were incubated overnight at 4°C in primary antibody solution, diluted in antibody dilutant (0.1% Triton-X, 0.5% NGS in PBS), washed in PBS, and incubated at RT for 1 hr in secondary antibody mixes diluted in PBS before mounting with Fluoromount-G. Antibodies used in this study are listed in the Antibodies table. Images were captured with a 63x oil immersion objective on either the LSM-900 laser scanning microscope with Airyscan (Zeiss) or Axio Imager Z2 fluorescence microscope equipped with Apotome (Zeiss). Final images are represented as maximum projection of Z-stacks.

### Protein lysis of retinal organoids

For bulk proteomic analysis of retinal organoids through development, 8-12 control and INPP5E^KD^ organoids were collected at d0 (undifferentiated iPSCs), d35 (early retinal organoid development), and d230 (mature retinal organoids). Organoids were washed two times in PBS and homogenized using the TissueLyser III (Qiagen) and lysed in RIPA buffer (50 mM Tris-HCl pH 7.5, 150 mM NaCl, 1% NP-40, 0.1% SDS (Life Technologies, 15553-027), 0.5% Sodium Deoxycholate, 1 mM UltraPure EDTA pH 8.0 (Invitrogen, 11568896)) containing cOmplete protease inhibitor cocktail (Roche, 11836170001). Samples were centrifuged for 15 mins at 14,000 rpm at 4°C. Protein concentration of the lysates was measured using the Pierce BCA protein assay (Life Technologies, 23225), and two technical replicates of 15 µg aliquots were snap frozen and stored at −80°C until mass spectrometry analysis.

### Liquid Chromatography-Mass spectrometry analysis and data analysis

Proteomic analysis was performed on retinal organoid samples as described previously (27,28). Briefly, tryptic peptides were processed for LC-MS/MS using data-independent acquisition mode (DIA) on the QExactive Plusmass spectrometer (Thermo Fisher Scientific). Raw data files were analyzed using DIA-NN 1.8.2 (29) in library-free mode against the human database (Swiss-Prot, May 2023, #20,401 proteins), and high-accuracy spectra with a minimum FDR of 0.01 and tryptic peptides were used for protein abundance label free quantification.

Statistical analysis was calculated using Perseus (version 1.6.15.0) (30). Significant proteins were identified using a Student’s T-test with permutation-based FDR of <0.05 and a Significance A (SignA) outlier test with Benjamini-Hochberg FDR correction of <0.05. Gene Ontology (GO) analysis was performed on significant proteins using the Gene Ontology Resource (31–33) for biological processes (BP), molecular functions (MF) and cellular component (CC). Enriched terms were filtered using a Fisher’s Exact test and corrected using the Bonferroni correction for multiple testing.

### Quantification and statistical analysis

Immunofluorescence images were quantified using Fiji software (34). For each timepoint analyzed, three to four organoids over two differentiations were analyzed. At least two images were taken per organoid per timepoint, and quantification was performed manually for area of expression, mean fluorescence intensity (MFI), length, or frequency of the relevant antibody staining. Statistical analyses were performed using GraphPad Prism 10 (GraphPad, San Diego, CA, USA). Comparison between groups was performed using either a Student’s *t* test or Mann-Whitney test, as indicated. Significance between groups was determined if *p* values were less than 0.05, and significance is indicated with asterisks (*, *p* =0.01-0.05; **, *p* =0.01-0.001; ***, *p*=0.001-0.0001; ****, *p*<0.0001).

## Data availability

All relevant data can be accessed in the article and supplementary material.

## Acknowledgements

We are grateful to Dr. Thomas Theil (University of Edinburgh, UK) for providing iPSC cell lines, the Radboudumc Technology Center Microscopy for use of their microscope facilities, Bernhard Schermer for providing the NPHP1 antibody, and Franziska Klose and Mohamed-Ali Jarboui from the Core Facility for Medical Proteomics (Eberhard Karls University of Tübingen, Germany). This work was supported by the European Union’s Horizon 2020 research and innovation program under the Marie Skłodowska-Curie grant agreement No. 861329 (‘SCilS’) to K.B. and R.R., by funding from the Dutch ministry of Education, Culture and Sciences, Gravitation grant: 024.006.034 Lifelong VISION to R.R. and by the EJP RD JTC 2022/ZonMW program ‘PREDICT’ (04630042210010).

## Author contributions

K.R.W. performed the experiments, analyzed the data and wrote the paper. M.A analyzed the data and wrote the paper. T.B, K.D. and K.B. performed and analyzed the M.S. experiments, R.R. coordinated the project, acquired funding and wrote the paper. All authors reviewed and edited the paper.

