## Supplemental Figures for "Loss of INPP5E affects photoreceptor outer segment membrane biogenesis in iPSC-derived human retinal organoids"

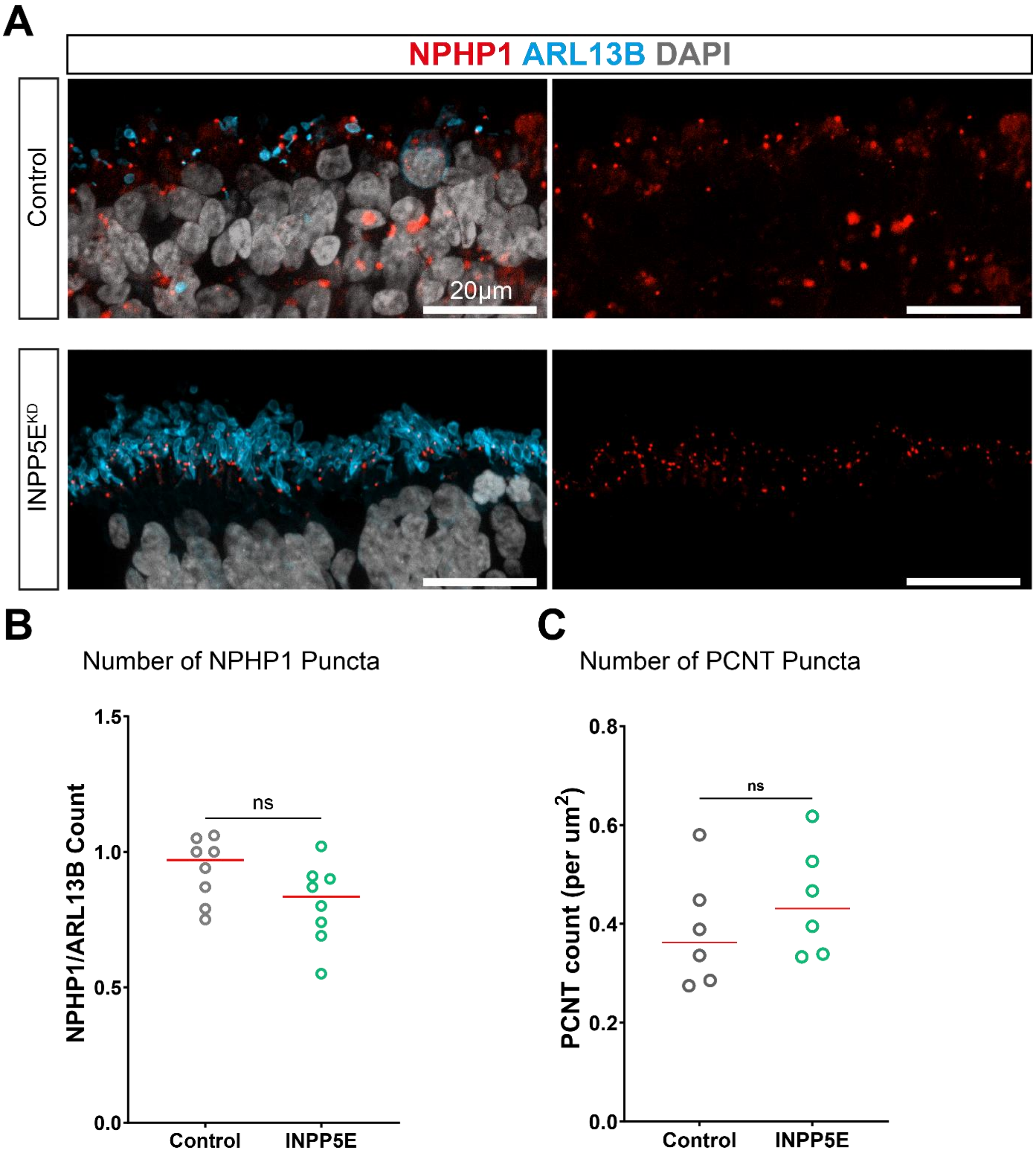


**Supplemental figure 1 – Basal body and transition zone markers are not affected in INPP5E$^{KD}$ organoids.** (A) Representative images of d230 ROs stained for NPHP1 (red), ARL13B (blue), and DAPI (grey). (B) Quantification of NPHP1 puncta as the count per ARL13B$^+$ cilia. (C) Pericentrin (PCNT) puncta count as the number of PCNT puncta per µm$^2$.

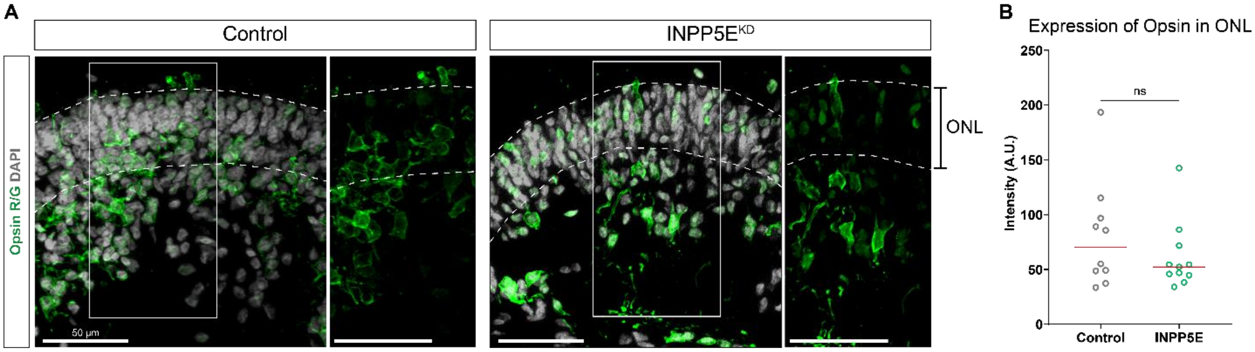


**Supplemental Figure 2 – Opsin localization is not altered in INPP5E$^{KD}$ ROs.** (A) Representative images of Control and INPP5E$^{KD}$ ROs stained for Opsin R/G (green) and DAPI (grey). Dashed lines represent the boundary of the ONL within the ROs (scale: 50 μm) (B) Quantification of Opsin expression in the ONL was measured as normalized intensity (A.U.)

Antibodies Table

| Antibody | Source | Identifier | Dilution |
|---|---|---|---|
| Mouse anti-CRX | Abnova | H00001406-M02 | 1:200 |
| Mouse anti-ARL13B (IgG2a) | NeuroMab | 75-287 | 1:500 |
| Rabbit anti-ARL13B | Proteintech | 17711-1-AP | 1:500 |
| Rabbit anti-Opsin R/G | Millipore | AB5405 | 1:200 |
| Mouse anti-Rhodopsin | Novus Biologicals | NBP2-59690-25ug | 1:500 |
| Rabbit anti-RCVN | Chemicon | AB5585 | 1:1000 |
| Mouse anti-PCNT (IgG1) | Abcam | AB28144 | 1:500 |
| Rabbit anti-INPP5E | Proteintech | 17797-1-AP | 1:500 |
| Rabbit anti-PCARE | Homemade | N/A | 1:250 |
| Mouse anti-GT335 | AdipoGen Life Sciences | AG-20B-0020-C100 | 1:250 |
| Mouse anti-NPHP1 | Bernhard Schermer Lab | N/A | 1:500 |
| Anti- Lectin PNA (568 conjugate) | Thermo Fisher Scientific | L32458 | 1:500 |
| Anti-Phalloidin (568 conjugate) | Invitrogen | A12380 | 1:250 |
| Goat anti-Mouse IgG1 (647) | Invitrogen | 10789384 | 1:500 |
| Goat anti-Mouse IgG1 (488) | Invitrogen | A21121 | 1:500 |
| Goat anti-Mouse IgG2a (647) | Thermo Fisher Scientific | A21241 | 1:500 |
| Goat anti-Mouse IgG2a (488) | Thermo Fisher Scientific | A21131 | 1:500 |
| Goat anti-Mouse IgG2a (568) | Invitrogen | A21134 | 1:500 |
| Goat anti-Mouse (488) | Invitrogen | A11029 | 1:500 |
| Goat anti-Mouse (568) | Life Technologies | A11031 | 1:500 |
| Goat anti-Rabbit (647) | Invitrogen | A21245 | 1:500 |
| Goat anti-Rabbit (488) | Molecular Probes | A11008 | 1:500 |
| Goat anti-Rabbit (568) | Molecular Probes | A11011 | 1:500 |
| DAPI | Thermo Fisher Scientific | D1306 | 1:100 |
